# Genetically Increased Telomere Length and Aging-related Physical and Cognitive Traits in the UK Biobank

**DOI:** 10.1101/538124

**Authors:** Kathryn Demanelis, Lin Tong, Brandon L. Pierce

**Affiliations:** Department of Public Health Sciences, University of Chicago, Chicago, IL, USA; Department of Human Genetics, University of Chicago, Chicago, IL, USA; University of Chicago Comprehensive Cancer Center, Chicago, IL, USA

**Keywords:** Telomeres, aging, UK Biobank, Mendelian randomization, blood pressure, pulmonary function, polygenic scores

## Abstract

**Background:** Telomere length (TL) shortens over time in most human cell types and is a potential biomarker aging. However, the causal impact of TL on physical and cognitive phenotypes that decline with age has not been extensively examined. Using a Mendelian randomization (MR) approach, we utilized genetically increased TL (GI-TL) to estimate the impact of TL on aging-related traits among UK Biobank (UKB) participants.

**Methods:** We manually curated >50 aging-related traits from UKB and restricted to unrelated participants of British ancestry (n=337,522). We estimated GI-TL as a linear combination of nine TL-associated SNPs, each weighted by its previously-reported association with leukocyte TL. Regression models were used to assess the associations between GI-TL and each trait. We obtained MR estimates using the two-sample inverse variance weighted (IVW) approach.

**Results:** We identified 5 age-related traits associated with GI-TL (Bonferroni-corrected threshold p<0.001): pulse pressure (PP) (p=5.2×10^−14^), systolic blood pressure (SBP) (p=2.9×10^−15^), diastolic blood pressure (DBP) (p=5.5×10^−6^), forced expiratory volume (FEV1) (p= p=0.0001), and forced vital capacity (FVC) (p=3.8×10^−6^). Under MR assumptions, one standard deviation increase in TL (∼1200 base pairs) increased PP, SBP, and DBP by 1.5, 2.3, and 0.8 mmHg, respectively, while FEV1 and FVC increased by 34.7 and 52.2 mL, respectively. The observed associations appear unlikely to be due to selection bias based on analyses including inverse probability weights and analyses of simulated data.

**Conclusions:** These findings suggest that longer TL increases pulmonary function and blood pressure traits. Further research is necessary to evaluate TL in cardiovascular and pulmonary age-related decline.

**KEY MESSAGES:**

- Telomere length (TL) is a potential biomarker and cause of aging, however, the causal relationship between TL and aging-related traits has not been thoroughly examined using a Mendelian randomization (MR) approach.
- We evaluated genetically increased TL (GI-TL) and its association with over 50 aging-related traits in the UK Biobank cohort using regression models and MR approaches.
- Pulmonary function (FEV1 and FVC) and blood pressure (SBP, DBP, and PP) traits were positively associated with GI-TL in the expected and unexpected direction, respectively.
- Using inverse probability weights to account for the non-representativeness of the UKB, our observed associations for GI-TL with blood pressure traits and pulmonary function persisted.
- Using simulated data to examine study selection as a potential source of collider bias, we concluded that selection bias was unlikely to explain the observed associations.

## INTRODUCTION

Telomeres are DNA-protein complexes that protect the ends of chromosomes from degradation and fusion. The DNA component, a 6-nucleotide repeat sequence, shortens with each cell division.^1^ Thus, telomere length (TL) decreases as human age in most cell types^2^ and is a proposed biomarker and potential cause of biologic aging.^3^ Meta-analyses of observational studies of leukocyte TL suggest that TL is associated with all-cause mortality,^4^ and risks for age-related chronic diseases including cardiovascular disease,^5^ type II diabetes,^6^ Alzheimer’s disease,^7^ and some cancers.^8^ These meta-analyses acknowledge that there is substantial heterogeneity across studies due to differences in TL measurement, study design, and adjustment for confounding factors, potentially affecting the validity of these associations.

Epidemiological studies of TL are susceptible to biases caused by confounding factors such as technical variation in TL measurement, environment, lifestyle, and cell type composition. Mendelian randomization (MR) is an alternative approach for estimating the impact of TL on disease susceptibility and utilizes genetic variants (SNPs) associated with leukocyte TL rather than measured TL itself, potentially avoiding biases caused by confounding factors and reverse causation.^9^ Providing certain assumptions are satisfied, SNPs associated with TL can be used to estimate the non-confounded causal relationship between TL and a health outcome. These assumptions include 1) the SNPs are associated with TL, 2) SNPs only affect the outcome via its effect on TL, and 3) SNPs are not associated with confounders that influence both TL and the outcome.^10^

SNPs associated with leukocyte TL have been identified in prior genome-wide association studies (GWAS).^11-13^ The ENGAGE consortium identified seven SNPs independently associated with TL, five of which were located in regions containing genes involved in telomere regulation and maintenance.^11^ Several prior studies have applied MR approaches by utilizing genetic determinants TL to assess the potential causal contribution of TL to overall and cause-specific mortality,^14^ the risk of cancer,^15^ and non-communicable diseases.^16^

While the biological basis for the role of TL in cellular aging is apparent, we do not know if variation in TL contributes to systemic aging of specific organ systems and the extent to which TL is causally related to age-related physical and cognitive decline. To address this question, we estimated genetically increased TL (GI-TL) and assessed its association with >50 aging-related traits in 337,522 unrelated UK Biobank (UKB) participants of British ancestry. Using a MR approach, we then estimated causal associations between TL and these traits using SNP summary statistics from the ENGAGE consortium and UKB cohort (**Figure 1**).

**Figure 1.**
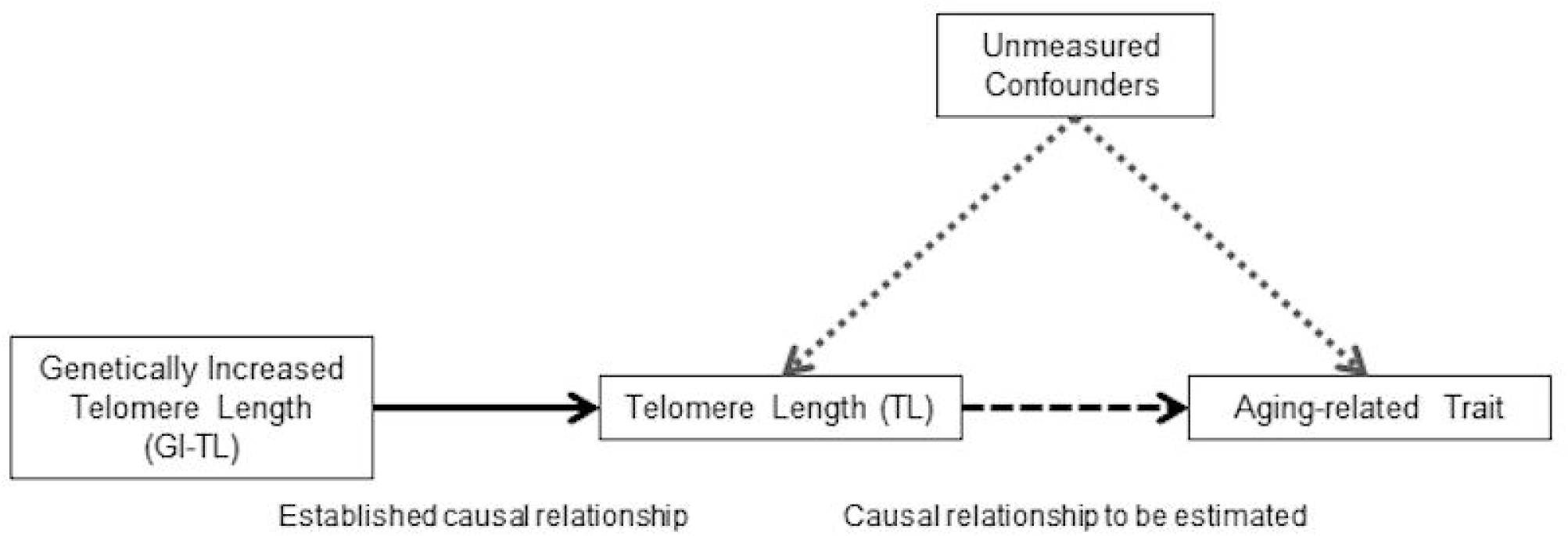
Causal diagram between telomere length (TL) and aging-related traits in UKB cohort.

## METHODS

### Study Sample

The UKB cohort is a prospective study of ∼500,000 middle-aged adults from the UK aged 40-69 years (recruited from 2006-2010). Postal invitations to participate were mailed to >9 million people registered with the UK National Health Service, who resided within 40 km of one of the 22 assessment centers.^17^ Of the adults invited, 5.45% responded and participated.^18^ At baseline, participants completed touchscreen questionnaires, physical exams, and verbal interviews, and a blood sample was obtained. All participants were genotyped at ∼800,000 SNPs on either the custom UK BiLEVE Axiom Array or UK Biobank Axiom Array, with the two arrays sharing 95% of SNPs. Further information related to array design, genotyping, imputation, and quality control is summarized elsewhere.^19^ We restricted our analysis to unrelated individuals with both self-reported and genetically predicted British ancestry based on the principal component (PC) analysis of genotypes. Our final sample size was 337,522 participants.

### Curation of Age-related Traits from UKB

We searched the UKB data showcase to manually curate a set of aging-related traits from the cognitive function, physical measures, and self-reported touchscreen questionnaire variables to capture the following domains of health (at baseline): cardiovascular, pulmonary, anthropometric, physical activity, sensory (hearing and eyesight), oral, musculoskeletal, pain and general health. Among the physical measurement variables, we extracted traits related to blood pressure (BP), arterial stiffness, spirometry, anthropometry, bone density, and hand grip strength. We utilized summarized cognitive function variables to capture traits related to visual memory, fluid intelligence, and short-term memory, as described in Bakrania et al.^20^ The remaining aging-related traits were extracted from the self-reported touchscreen questionnaire, and these traits included those related to pain, weight change, activity, fractures, oral health, hearing loss, eyesight, and general health. We extracted a total of 52 aging-related traits.

### Derivation of Frailty Index

Frailty indices (FI) have been proposed to capture the variation in health among aging individuals. We applied the algorithm proposed by Williams et al to compute a FI for each participant.^21^ Briefly, 49 self-reported questionnaire variables related to health, disease, disability, and mental wellbeing were extracted and coded into binary (0 or 1) or ordinal variables (values ranging from 0 to 1). For each UKB participant, FI was computed from the sum of the observations across these 49 variables and divided by 49 and expressed as a percent. For participants with missing data for one or more variables, values for missing observations were imputed using multiple imputation by chained equations.^22^

### Estimation of Genetically Increased TL (GI-TL)

We identified SNPs associated with leukocyte TL from prior genome-wide meta-analyses.^11-13^ We selected nine SNPs that were associated with leukocyte TL and not in linkage disequilibrium with each other (see **Table 1**). For each SNP, we extracted the association estimate (i.e., beta coefficient and standard error) corresponding to the impact of a one allele increase on leukocyte TL in terms of the standard deviation (SD) (∼1200 base pairs (bp)) from the ENGAGE telomere consortium.^11^ GI-TL was estimated as a weighted linear combination of the TL-associated SNPs, corresponding to the sum of the weighted values for each SNP 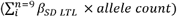. GI-TL was interpreted as the additional TL relative to an arbitrary TL value, corresponding to GI-TL=0.^23, 24^ A hypothetical individual with zero “long” TL alleles will have a GI-TL=0, corresponding to no additional bp of TL, while a hypothetical individual with 18 “long” TL alleles will have a GI-TL=1.08, corresponding to ∼1296 additional bp of TL (compared to an individuals with zero “long” alleles) (**Figure S1**).

### Statistical Analysis

Each aging-related trait was expressed as a binary or continuous variable. For individuals taking BP medication at baseline, we added 15 mmHg and 10 mmHg to their baseline systolic (SBP) and diastolic (DBP) BP, respectively,^25^ and averaged first and second measurements. Hypertension was defined as >140 mmHg SBP or >90 mmHg DBP or self-reported BP medication use. For continuous variables we removed outlying or implausible values for SBP, DBP, forced expiratory volume (FEV1), forced vital capacity (FVC), peak expiratory flow (PEF1), and pulse rate. We computed waist to hip ratio from waist and hip measurements, pulse pressure (PP) from the difference between SBP and DBP, and FV-ratio of FEV1 to FVC. We computed genotyping PCs among the selected UKB participants using EIGENSOFT^26-28^ implemented in PLINK 1.9.

Logistic or linear multivariable regression models were utilized to estimate associations for each aging-related trait with GI-TL. We conducted analyses stratified by gender and median age (< or ≥ 58 years). A Bonferroni correction was applied to account for multiple testing at α=0.05. We utilized Cox proportional hazards models to estimate the association between GI-TL and mortality. Follow-up began when the participant conducted their baseline assessment and ended at whichever happened first: death, departure from study, or censoring date (November 30, 2015). All analyses included the following covariates: age, sex, and ten genotyping PCs.

### Mendelian Randomization Analysis

We used a two-sample inverse-weighted variance (IVW) approach to obtain estimates of the effect of TL on aging-related traits, as described previously. ^9, 29^ For each SNP included in the GI-TL score (k=9 SNPs), the beta coefficient (X_k_) and its standard error (se) were obtained from the summary statistics from the GWAS of leukocyte TL from the ENGAGE consortium, and the per allele association estimate (Y_k_) and its se 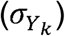 with the aging-related trait were obtained from the UKB cohort. The IVW MR estimate and its se were computed as follows:

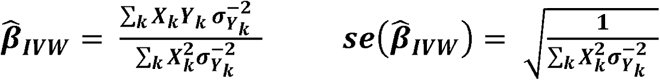

All MR analyses were executed using the *MendelianRandomization* package in R.^30^ We compared the MR-estimates from IVW to MR-estimates obtained from the maximum likelihood^9, 31^ and weighted median^32^ approaches to evaluate the consistency of the MR estimates. From MR-Egger regression, we extracted intercept-test for directional pleiotropy to evaluate potential violation of MR assumptions.^32, 33^ To examine horizontal pleiotropic outliers, we applied the recently developed Mendelian Randomization Peliotropy RESidual Sum and Ouliter (MR-PRESSO) test.^34^

### Inverse Probability Weighting

Selection can be a form of collider bias in MR studies when confounders are present. Selection into the study based on the outcome or/and the risk factor of interest can bias the MR effect estimate (**Figure S2**), potentially violating the assumption that the instrumental variable and confounder(s) are marginally independent.^35, 36^ UKB participants are more likely to be female, healthier, older, and of higher social and economic status than the target source population.^18^ We applied an inverse probability weighting (IPW) approach to create a pseudo-population that is unaffected by selection on factors like age, sex, smoking status, and BP,^37^ in order to estimate the average causal effect of TL (on BP and pulmonary traits) in the general population.^36^ We computed IPWs as follows, using the percent of the source population belonging to a particular group based on a set of characteristics (up to four levels: i,j,k,l) and the percent of UKB participants belonging to that group:

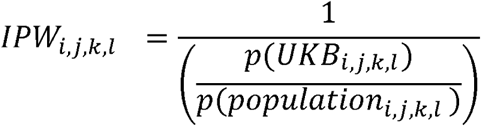

Using Web Table 1 in Fry et al,^18^ we extracted information on the gender and age group distribution (40-50, 50-60, 60-70 years) of the UKB source population. Because there were no cross-tabulated data available for the UKB source population, we utilized the Health Survey of England (HSE) 2008 and extracted sets of population-weighted cross-tabulated proportions for groups based on sex, age (40-50, 50-60, 60-70 years), smoking status, and hypertension from the HSE.^38^ We fit linear models adjusted for age, sex and 10 genotyping PCs and included IPWs as weights.

### Simulated Datasets to Examine the Impact of Selection Bias on TL and SBP

We conducted simulations to examine how selection on TL, SBP (the strongest association identified), and/or confounders influenced the MR estimates under the hypothesis of no causal effect (β_x_=0) and of a true effect (based on the observed association) (β_x_=0.12). Using an approach similar to that described by Gkatzionis and Burgess,^36^ we generated the risk factor, TL, (X_i_) as a linear combination of a genetic risk score, a confounder, and an independent error term (see equation 1), where the genetic risk score, confounder and random error term are independently drawn from a N(0,1) distribution. The non-varying parameters included percent variation explained (PVE) by genetic instrument for TL (α^2^_g_=0.04), the estimated amount of variation explained by SNPs associated with TL, and PVE in TL by the confounder that we assumed (α^2^_u_=0.5).

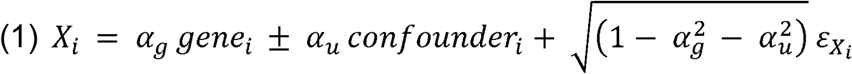

We generated the outcome, SBP, using a linear combination of the risk factor (X_i_), generated in equation 1, confounder, and an independent error term (see equation 2), where the random error term is drawn from N(0,1) distribution. The non-varying parameters included PVE explained by confounders in SBP that we assumed (β^2^_u_=0.50), and the causal effect to be estimated (β_x_=0 or β_x_=0.12).

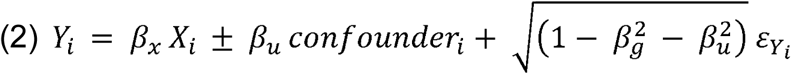

For each simulated observation, a selection probability was generated and assigned 0 (not selected) or 1 (selected) based on a Bernoulli trial with that selection probability. We examined two functions to generate the selection probability: the risk factor effect on selection (see equation 3) and the outcome effect on selection (see equation 5). We varied the following parameters within the selection probability: the effect of risk factor on selection (γ_x_), the effect of outcome on selection (γ_y_), and the confounder effect on selection (γ_u_) (see **Appendix S1**). We also included the baseline prevalence of selection into the UKB (γ_0_=0.0545).

Risk factor effect on selection:

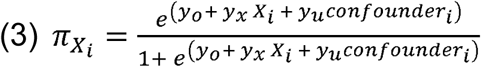

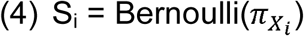

Outcome effect on selection:

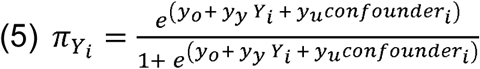

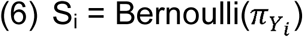

For each simulated dataset, we generated a random sample of 10,000,000 observations, similar to the size of the UKB source population. Among the observations with S=1, we randomly selected 500,000 observations. To obtain the causal effect estimate of the risk factor on the outcome, we conducted linear regressions of the genetic instrument (gene) on the risk factor (X) and the genetic instrument (gene) on the outcome (Y), and then, computed the ratio estimate and its se. We ran 1000 simulations per scenario and extracted the mean, median, SD, median se of the ratio estimates.

All analyses were performed in R 3.4.3.

## RESULTS

We analyzed 52 aging-related traits among the 337,522 UKB participants. The median age of participants included was 58 years (range: 39 to 72 years), and there were more women (53.7%) than men. Fifteen aging-related traits were associated with GI-TL at a significance threshold p=0.05 (**Table 2**), but only five traits surpassed the Bonferroni-corrected significance threshold (p<0.001) (**Figure 2**). GI-TL was associated with better pulmonary function. FEV1 and FVC increased by 34.7 mL (95% CI [16.9, 52.6], p=0.0001) and 52.2 mL (95% CI [30.1, 74.4], p=3.8×10^−6^) per one SD increase in TL (corresponding to ∼1,200 bp), respectively. The association between GI-TL and FVC was stronger among men (β=79.4 mL, 95% CI [41.1, 117.6], p=4.8×10^−5^) compared to women (β=29.4 mL, 95% CI [4.5, 54.3], p=0.02) (interaction p=0.02) (**Table S1** and **Figure S3**). When stratified by median age, GI-TL was positively associated with FEV1 (β=49.7 mL, 95% CI [25.5, 73.8], p=5.7×10^−5^) and FVC (β=69.9 mL, 95% CI [40.1, 99.7], p=4.3×10^−6^) among individuals aged ≥ 58 years but FEV1 and FVC were not associated with GI-TL among individuals younger than 58 years (p>0.05) (**Table S2** and **Figure S4**), with interaction p-values of 0.05 and 0.07, respectively.

**Figure 2.**
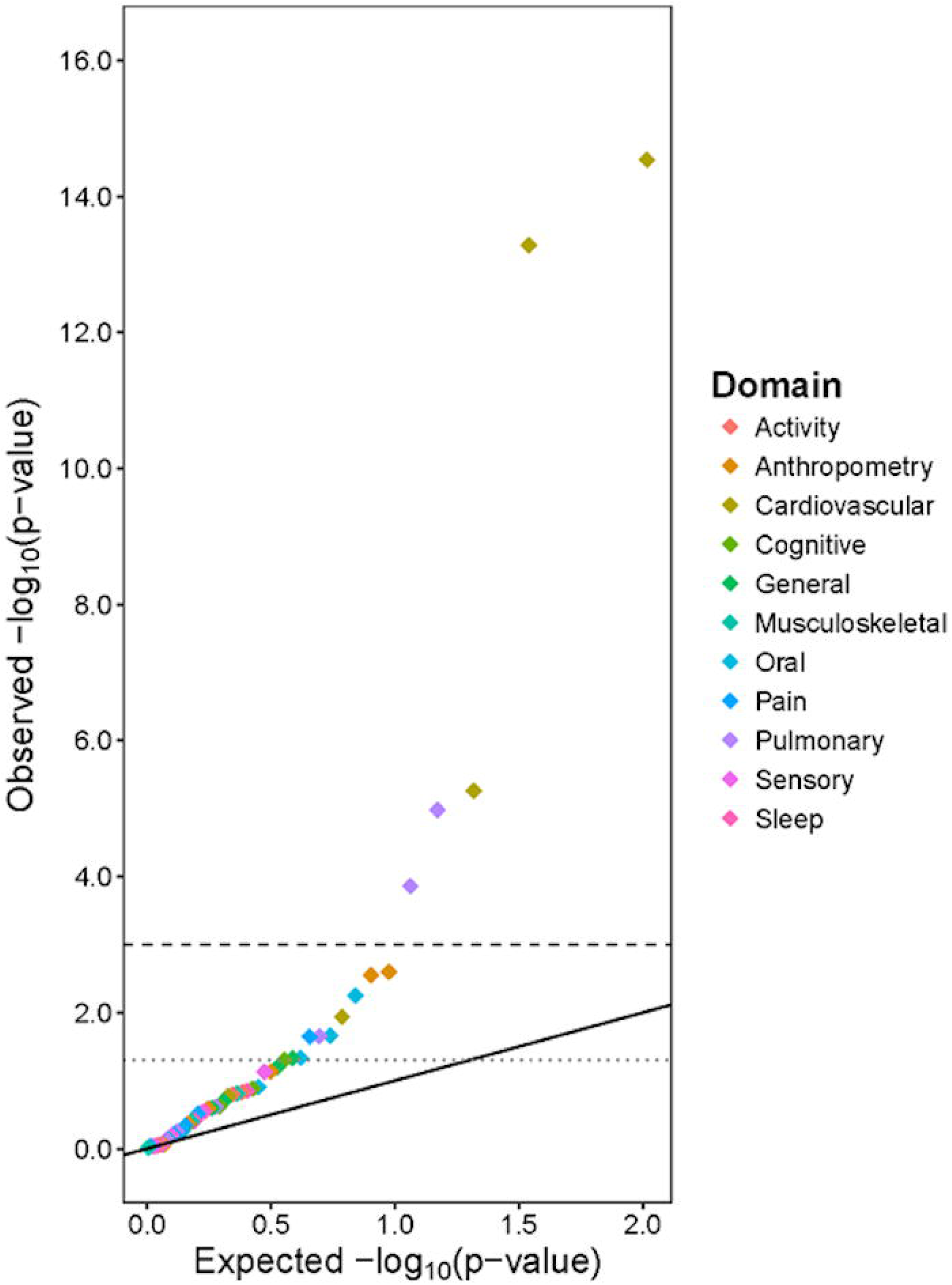
Quantile-quantile plot of genetically increased telomere length (GI-TL) and age-related traits among UK Biobank participants.

Half of the UKB participants were defined as hypertensive (50.1%), and age was correlated with SBP (Pearson’s r=0.332), DBP (r=0.071), and PP (r=0.418). We observed that GI-TL was associated with increased SBP (β=2.3 mmHg, 95% CI [1.7, 2.9], p=2.9×10^−15^), DBP (β=0.8 mmHg, 95% CI: [0.4, 1.1], p=5.5×10^−6^), and PP (β=1.5 mmHg, 95% CI [1.1, 1.9], p=5.2×10^−14^) per one SD increase in TL. GI-TL was positively associated with SBP, PP, and DBP among both men and women, and there was not strong evidence that these associations differed by sex (**Table S1**). When we stratified by median age (< or >= 58 years), GI-TL was positively associated with SBP, PP, and DBP within both strata (**Table S2**).

Among our study participants, 9196 deaths occurred before December 2015, and GI-TL was not associated with overall mortality, death from ischemic heart disease (n=1053), cerebrovascular disease (n=327), or chronic lower respiratory disease (n=227) (**Table S3**). GI-TL was associated with decreased risk of mortality from idiopathic pulmonary fibrosis (IPF) (HR=0.10, 95% CI: [0.02, 0.55], p=0.008, n=106). The *TERT* long allele (rs2736100) was associated with increased cerebrovascular disease mortality risk (HR=1.20, 95% CI [1.03, 1.40], p=0.02) and with decreased IPF risk (HR=0.75, 95% CI [0.57, 0.99], p=0.04).

In an exploratory sensitivity analysis, we sought to examine whether our results were influenced by the selection bias related to participation in the UKB cohort. To do this, we applied IPWs in linear regression models to examine the association between GI-TL and the five significant aging-related traits (see **Methods**). When we applied IPWs based on age only (derived from the UKB source population), we observed that the associations were slightly attenuated but still highly significant between GI-TL and SBP (β=2.2, 95% CI [1.6, 2.7], p=1.7×10^−14^) and PP (β=1.3, 95% CI [1.0, 1.7], p=4.8×10^−13^) (**Table S4**). When we applied IPWs based on age, sex, and hypertension status (derived from the HSE), the association between GI-TL and SBP and PP were attenuated by 13.1% (β=2.0, 95% CI [1.5, 2.5], p=1.7×10^−13^) and 17.3% (β=1.2, 95% CI [0.9, 1.6], p=4.5×10^−12^), respectively. The associations between FVC and GI-TL were strengthened when IPWs were applied. Overall, the results we observed persisted after the inclusion of IPWs to account for underlying factors affecting selection into and participation in the UKB.

These pulmonary and BP traits were further examined using a two-sample MR analysis. Provided the MR assumptions are satisfied,^39^ MR-estimates can be interpreted as the effect of a one SD increase in TL (∼1,200 bp) on each trait.^11^ Using the IVW MR method, FEV1 and FVC increased by 34.7 mL (95% CI [7.9, 61.5], p=0.01) and 52.2 mL (95% CI [18.2, 86.2], p=0.003) per one SD increase in TL, respectively (**Figure 3 and Table S5**). For three out of the nine SNPs, the “long TL” allele was positively associated with FVC: *TERC* (β=9.2 mL, 95% CI [5.1,13.3], p=1.2×10^−5^), *TERT* (β=4.7 mL, 95% CI [1.1, 8.2], p=0.01) and *RTEL1* (β=8.7 mL, 95% CI [3.5,13.9], p=0.001). For FEV1, per-allele associations were observed for *TERC* (β=6.4 mL, 95% CI [0.0, 8.2], p=1.4×10^−4^), *RTEL1* (β=5.4 mL, 95% CI [1.2, 9.6], p=0.01), and *ACYP2* (β=4.1 mL, 95% CI [5.1,13.3], p=0.05).

**Figure 3.**
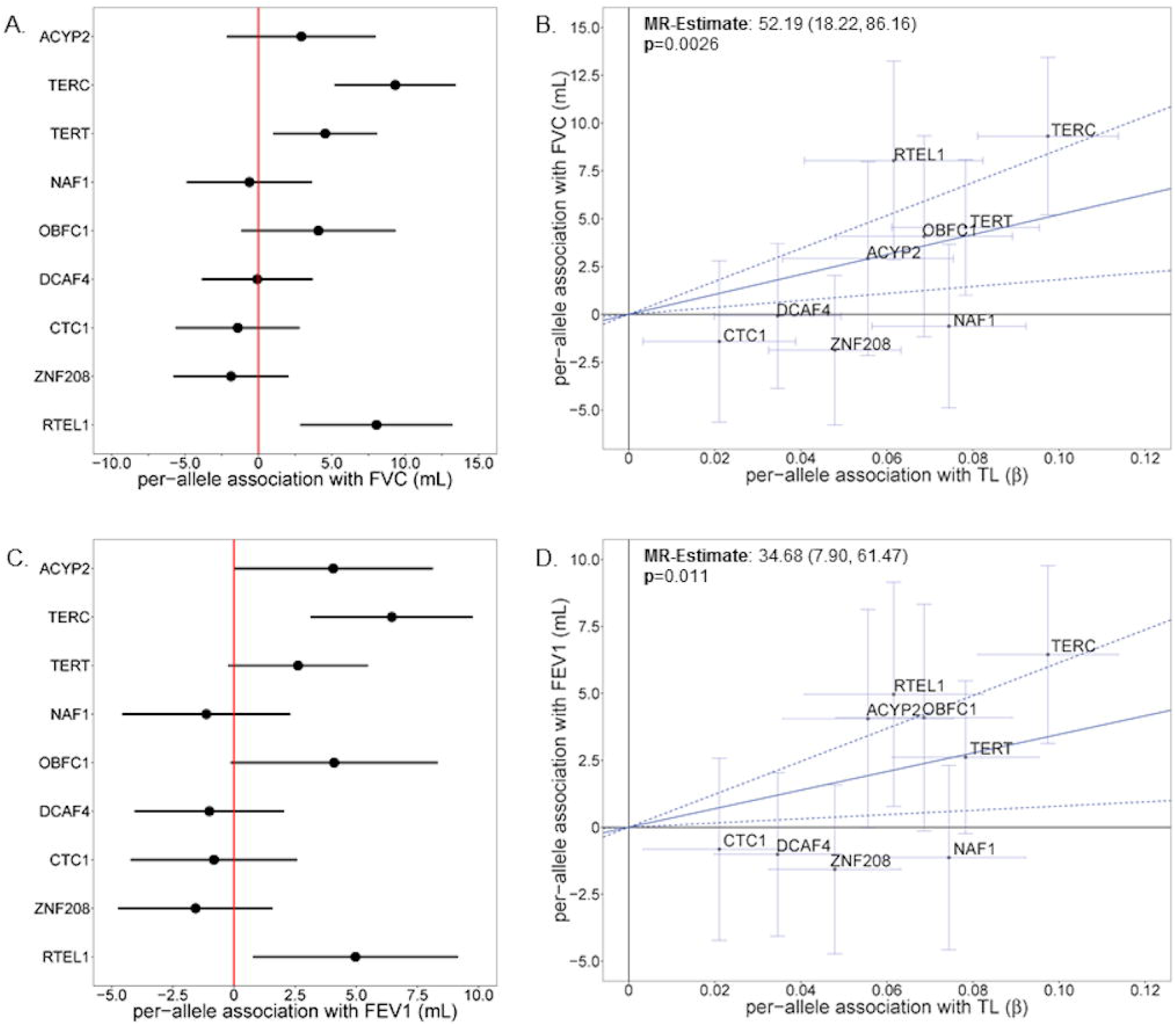
Long TL alleles increased pulmonary function traits.

MR analysis indicated that a one SD increase in TL increased SBP, DBP, and PP by 2.3 mmHg (95% CI [1.3, 3.3], p=8.2×10^−6^), 0.8 mmHg (95% CI [0.3, 1.2], p=0.001), and 1.5 mmHg (95% CI [0.5, 2.5], p=0.002), respectively (**Figure 4**). SBP was positively associated with eight of the nine TL-associated SNPs while DBP was only positively associated with three of the nine SNPs (p<0.05). DBP was not strongly associated with either the *TERT* or *TERC* SNPs (p>0.05). Six of the nine TL-associated SNPs were positively associated with PP (p<0.05).

**Figure 4.**
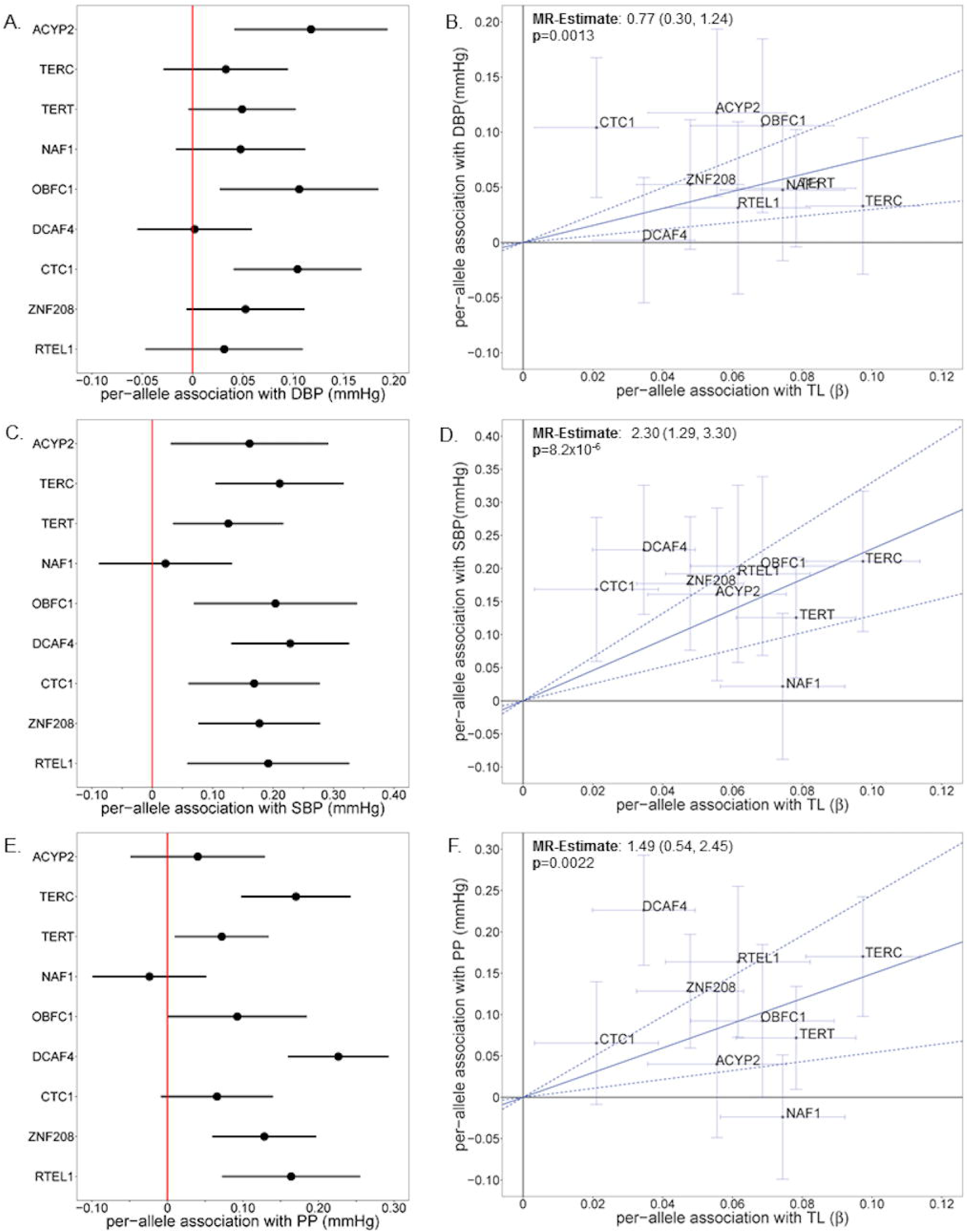
Long TL alleles increased blood pressure traits.

In addition to the IVW MR method, we also obtained MR estimates using several alternative MR approaches (**Table S5**). For both BP and the pulmonary traits, the maximum likelihood approach yielded similar MR-estimates and p-values. Using MR-Egger, the estimated intercept differed from zero for SBP (p=0.001), suggesting potential bias in the MR-estimate. However, the MR-estimates obtained from the weighted median approach were similar to IVW for all traits. When we applied the MR-PRESSO test to identify horizontal pleiotropic outliers, NAF1 and DCAF4 were identified as outliers for SBP and PP (p<0.05), however, the removal the outliers increased the IVW MR-estimates for both traits.

In datasets simulated under the null hypothesis of no causal effect (β=0), selection on a continuous outcome (such as SBP) did not introduce bias or affect the false positive rate (**Table S6**). However, strong selection on a risk factor (e.g. TL) increased the false positive rate and biased the MR-estimate away from the null, with the direction of the bias depending on the direction of the confounder effects (**Table S7**). Notably, this selection would have to be extremely strong (e.g. y_x_=-2 or y_x_=2) in order to generate the association of the magnitude we observe between TL and SBP (β_x_=0.12). Under the alternative hypothesis of a true effect between SBP and TL (β_x_=0.12), selection on the outcome attenuated causal effect estimate but did not decrease the empirical power to detect a true association of this magnitude between SBP and TL (**Table S8**). When there was selection on the risk factor (again under the alternative hypothesis with β_x_=0.12), two different scenarios were observed (**Table S9**). First, if the confounder effect on the risk factor and the outcome was in the same direction, then selection on the risk factor attenuated the association between SBP and TL and decreased the empirical power when there was also selection on the confounder (y_u_=-1 or y_u_=1). Second, if the confounder effect on the risk factor and outcome were in opposite directions, then strong selection on the risk factor biased the association between SBP and TL upward (similar in nature to the bias observed in **Table S7**).

## DISCUSSION

In this work, we examined the association between GI-TL and >50 aging-related traits among the UKB participants of British ancestry. While most aging-related traits did not show clear associations with GI-TL, we observed positive associations for GI-TL with pulmonary function and BP traits. MR results suggest that longer TL is causally related to increased BP (SBP, DBP, and PP) and pulmonary function (FEV1 and FVC). Additionally, the associations of GI-TL with FEV1 and FVC were stronger among older UKB participants, suggesting longer TL may protect against age-related pulmonary decline.

Compared to our results, prior studies have observed similar associations between TL and pulmonary function. Among 46,396 Danish individuals, higher FEV1 and FVC was associated with longer measured TL.^40^ A meta-analysis of 14 European studies identified positive associations between measured TL and FEV1 and FVC.^41^ While FEV1 and FVC have not been examined in MR studies of TL, a large MR study of TL found evidence for an effect of long TL on decreased risk of interstitial lung disease (similar to IPF).^16^ IPF is a hallmark of telomeropathy syndromes, characterized by very short TL due to mutations in telomere maintenance genes, such as TERT and TERC.^42, 43^ The relationship between TL and pulmonary diseases indicates that TL may be a biologically important factor that contributes to the age-related decline of pulmonary function.

Prior studies have observed inconsistent associations between TL and BP traits, with several null results reported.^44-49^ SBP was positively associated with measured TL in adult NHANES participants with SBP above 140 mmHg,^50^ in Costa Rican elderly adults,^51^ and in Australian adults 40-44 years.^52^ DBP was inversely associated with measured TL in NHANES^50^ and Costa Rican elderly adults.^51^ In Danish population-based cohorts, longer measured TL was inversely associated with SBP while more TL “shortening” alleles were associated with decreased SBP.^14^ Correspondingly, the percent of individuals with hypertension decreased with increasing TL “shortening” alleles.^53^ The opposite associations between GI-TL and measured TL for BP traits suggest that these potential differences may be due to residual confounding (due to biologic, technical, and/or environmental effects) and/or biologic pleiotropy related to GI-TL.

Our analysis has several strengths and limitations. The UKB sample size is very large, which is critical for precise MR-based estimation. UKB also has a uniform and rigorously collected set of aging-related traits. GI-TL is constructed based on several SNPs that are near genes with clear relevance to TL biology, making GI-TL an instrumental variable with strong biological connection to TL. However, a central limitation of MR is that it is not possible to completely rule out the possibility of biologic pleiotropy between our TL-associated SNPs and aging-related traits. An additional limitation is the non-representativeness of the UKB sample, potentially reducing the external validity of our results; this limitation we attempted to address using simulated data and IPWs. We utilized IPWs to create pseudo-populations that are unaffected by selection on certain factors and more closely resemble the UKB source population. While all association estimates remained highly significant after the inclusion of IPWs, the association between GI-TL and BP traits slightly decreased and the association between GI-TL and pulmonary traits increased, suggesting that IPWs should be further explored for other characteristics that may influence participation (e.g. pulmonary function) and further developed for large cohorts such as UKB to enhance the validity and reliability of MR estimates. From our analysis of simulated data to examine selection as a source of collider bias, we concluded that neither selection on the outcome (e.g. SBP) nor on the risk factor (e.g. TL) were likely to explain our observed MR association between TL and SBP. While we cannot create IPWs based on TL, selection on TL would have to be very strong to produce the observed association (under the null hypothesis of no effect), which in our view is not a biologically plausible explanation for the observed associations. These results based on simulated data were consistent with what was observed in the simulations reported by Gkatzionis and Burgess.^36^

In conclusion, among unrelated UKB participants of British ancestry, GI-TL was associated with increased pulmonary function but also with higher BP. These findings provide evidence for a causal relationship between TL and these aging-related traits. Future longitudinal studies should evaluate age-related temporal changes in pulmonary function and BP traits and how these related to TL.

## ACKNOWLEDGEMENTS

This work was supported by active and past National Institute of Health grants [R35ES028379, R01ES020506, and U01HG007601 to B.L.P]. We acknowledge fellowship support for Dr. Demanelis provided by the National Institute of Aging Specialized Demography and Economics of Aging Training Program and NIH Research Supplement to Promote Diversity in Health-Related Research [T32AG000243 and R35ES028379-02S1 to K.D.] at the University of Chicago. We would like to thank the UK Biobank leadership, staff, and participants for their contributions to this research and cohort. Figure 1. Causal diagram between telornere length (TL) and aging-related traits in Ul<B cohort.

## REFERENCES

1. d’Adda di Fagagna F. Living on a break: cellular senescence as a DNA-damage response. Nat Rev Cancer 2008; 8: 512–22.

2. Daniali L, Benetos A, Susser E, et al. Telomeres shorten at equivalent rates in somatic tissues of adults. Nat Commun 2013; 4: 1597.

3. Blackburn EH, Epel ES, Lin J. Human telomere biology: A contributory and interactive factor in aging, disease risks, and protection. Science 2015; 350: 1193–8.

4. Mons U, Muezzinler A, Schottker B, et al. Leukocyte Telomere Length and All-Cause, Cardiovascular Disease, and Cancer Mortality: Results From Individual-Participant-Data Meta-Analysis of 2 Large Prospective Cohort Studies. Am J Epidemiol 2017; 185: 1317–26.

5. Haycock PC, Heydon EE, Kaptoge S, Butterworth AS, Thompson A, Willeit P. Leucocyte telomere length and risk of cardiovascular disease: systematic review and meta-analysis. BMJ 2014; 349: g4227.

6. Willeit P, Raschenberger J, Heydon EE, et al. Leucocyte telomere length and risk of type 2 diabetes mellitus: new prospective cohort study and literature-based meta-analysis. PLoS One 2014; 9: e112483.

7. Forero DA, Gonzalez-Giraldo Y, Lopez-Quintero C, Castro-Vega LJ, Barreto GE, Perry G. Meta-analysis of Telomere Length in Alzheimer’s Disease. J Gerontol A Biol Sci Med Sci 2016; 71: 1069–73.

8. Zhang X, Zhao Q, Zhu W, et al. The Association of Telomere Length in Peripheral Blood Cells with Cancer Risk: A Systematic Review and Meta-analysis of Prospective Studies. Cancer Epidemiol Biomarkers Prev 2017; 26: 1381–90.

9. Burgess S, Butterworth A, Thompson SG. Mendelian randomization analysis with multiple genetic variants using summarized data. Genet Epidemiol 2013; 37: 658–65.

10. Boef AG, Dekkers OM, le Cessie S. Mendelian randomization studies: a review of the approaches used and the quality of reporting. Int J Epidemiol 2015; 44: 496–511.

11. Codd V, Nelson CP, Albrecht E, et al. Identification of seven loci affecting mean telomere length and their association with disease. Nat Genet 2013; 45: 422–7, 7e1-2.

12. Mangino M, Christiansen L, Stone R, et al. DCAF4, a novel gene associated with leucocyte telomere length. J Med Genet 2015; 52: 157–62.

13. Mangino M, Hwang SJ, Spector TD, et al. Genome-wide meta-analysis points to CTC1 and ZNF676 as genes regulating telomere homeostasis in humans. Hum Mol Genet 2012; 21: 5385–94.

14. Rode L, Nordestgaard BG, Bojesen SE. Peripheral blood leukocyte telomere length and mortality among 64,637 individuals from the general population. J Natl Cancer Inst 2015; 107: djv074.

15. Rode L, Nordestgaard BG, Bojesen SE. Long telomeres and cancer risk among 95 568 individuals from the general population. Int J Epidemiol 2016; 45: 1634–43.

16. Telomeres Mendelian Randomization C, Haycock PC, Burgess S, et al. Association Between Telomere Length and Risk of Cancer and Non-Neoplastic Diseases: A Mendelian Randomization Study. JAMA Oncol 2017; 3: 636–51.

17. Sudlow C, Gallacher J, Allen N, et al. UK biobank: an open access resource for identifying the causes of a wide range of complex diseases of middle and old age. PLoS Med 2015; 12: e1001779.

18. Fry A, Littlejohns TJ, Sudlow C, et al. Comparison of Sociodemographic and Health-Related Characteristics of UK Biobank Participants With Those of the General Population. Am J Epidemiol 2017; 186: 1026–34.

19. Bycroft C, Freeman C, Petkova D, et al. The UK Biobank resource with deep phenotyping and genomic data. Nature 2018; 562: 203–9.

20. Bakrania K, Edwardson CL, Khunti K, Bandelow S, Davies MJ, Yates T. Associations Between Sedentary Behaviors and Cognitive Function: Cross-Sectional and Prospective Findings From the UK Biobank. Am J Epidemiol 2018; 187: 441–54.

21. Williams DM, Jylhava J, Pedersen NL, Hagg S. A Frailty Index for UK Biobank Participants. J Gerontol A Biol Sci Med Sci 2018.

22. White IR, Royston P, Wood AM. Multiple imputation using chained equations: Issues and guidance for practice. Stat Med 2011; 30: 377–99.

23. Walsh KM, Codd V, Rice T, et al. Longer genotypically-estimated leukocyte telomere length is associated with increased adult glioma risk. Oncotarget 2015; 6: 42468–77.

24. Hamad R, Walter S, Rehkopf DH. Telomere length and health outcomes: A two-sample genetic instrumental variables analysis. Exp Gerontol 2016; 82: 88–94.

25. Warren HR, Evangelou E, Cabrera CP, et al. Genome-wide association analysis identifies novel blood pressure loci and offers biological insights into cardiovascular risk. Nat Genet 2017; 49: 403–15.

26. Price AL, Patterson NJ, Plenge RM, Weinblatt ME, Shadick NA, Reich D. Principal components analysis corrects for stratification in genome-wide association studies. Nat Genet 2006; 38: 904–9.

27. Galinsky KJ, Loh PR, Mallick S, Patterson NJ, Price AL. Population Structure of UK Biobank and Ancient Eurasians Reveals Adaptation at Genes Influencing Blood Pressure. Am J Hum Genet 2016; 99: 1130–9.

28. Galinsky KJ, Bhatia G, Loh PR, et al. Fast Principal-Component Analysis Reveals Convergent Evolution of ADH1B in Europe and East Asia. Am J Hum Genet 2016; 98: 456–72.

29. Zhang C, Doherty JA, Burgess S, et al. Genetic determinants of telomere length and risk of common cancers: a Mendelian randomization study. Hum Mol Genet 2015; 24: 5356–66.

30. Yavorska OO, Burgess S. MendelianRandomization: an R package for performing Mendelian randomization analyses using summarized data. Int J Epidemiol 2017; 46: 1734–9.

31. Thompson JR, Minelli C, Abrams KR, Tobin MD, Riley RD. Meta-analysis of genetic studies using Mendelian randomization--a multivariate approach. Stat Med 2005; 24: 2241–54.

32. Bowden J, Davey Smith G, Haycock PC, Burgess S. Consistent Estimation in Mendelian Randomization with Some Invalid Instruments Using a Weighted Median Estimator. Genet Epidemiol 2016; 40: 304–14.

33. Burgess S, Thompson SG. Interpreting findings from Mendelian randomization using the MR-Egger method. Eur J Epidemiol 2017; 32: 377–89.

34. Verbanck M, Chen CY, Neale B, Do R. Detection of widespread horizontal pleiotropy in causal relationships inferred from Mendelian randomization between complex traits and diseases. Nat Genet 2018; 50: 693–8.

35. Munafo MR, Tilling K, Taylor AE, Evans DM, Davey Smith G. Collider scope: when selection bias can substantially influence observed associations. Int J Epidemiol 2018; 47: 226–35.

36. Gkatzionis A, Burgess S. Contextualizing selection bias in Mendelian randomization: how bad is it likely to be? Int J Epidemiol 2018.

37. Nohr EA, Liew Z. How to investigate and adjust for selection bias in cohort studies. Acta Obstet Gynecol Scand 2018; 97: 407–16.

38. National Centre for Social Research UCL, Department of Epidemiology and Public Health. Health Survey for England, 2008. 4th Edition ed: UK Data Service; 2013.

39. Didelez V, Sheehan N. Mendelian randomization as an instrumental variable approach to causal inference. Stat Methods Med Res 2007; 16: 309–30.

40. Rode L, Bojesen SE, Weischer M, Vestbo J, Nordestgaard BG. Short telomere length, lung function and chronic obstructive pulmonary disease in 46,396 individuals. Thorax 2013; 68: 429–35.

41. Albrecht E, Sillanpaa E, Karrasch S, et al. Telomere length in circulating leukocytes is associated with lung function and disease. Eur Respir J 2014; 43: 983–92.

42. Armanios M, Blackburn EH. The telomere syndromes. Nat Rev Genet 2012; 13: 693–704.

43. Armanios M. Telomeres and age-related disease: how telomere biology informs clinical paradigms. J Clin Invest 2013; 123: 996–1002.

44. Bekaert S, De Meyer T, Rietzschel ER, et al. Telomere length and cardiovascular risk factors in a middle-aged population free of overt cardiovascular disease. Aging Cell 2007; 6: 639–47.

45. Brouilette SW, Moore JS, McMahon AD, et al. Telomere length, risk of coronary heart disease, and statin treatment in the West of Scotland Primary Prevention Study: a nested case-control study. The Lancet 2007; 369: 107–14.

46. Cheng YY, Kao TW, Chang YW, et al. Examining the gender difference in the association between metabolic syndrome and the mean leukocyte telomere length. PLoS One 2017; 12: e0180687.

47. Fitzpatrick AL, Kronmal RA, Gardner JP, et al. Leukocyte telomere length and cardiovascular disease in the cardiovascular health study. Am J Epidemiol 2007; 165: 14–21.

48. Denil SL, Rietzschel ER, De Buyzere ML, et al. On cross-sectional associations of leukocyte telomere length with cardiac systolic, diastolic and vascular function: the Asklepios study. PLoS One 2014; 9: e115071.

49. Revesz D, Milaneschi Y, Verhoeven JE, Lin J, Penninx BW. Longitudinal Associations Between Metabolic Syndrome Components and Telomere Shortening. J Clin Endocrinol Metab 2015; 100: 3050–9.

50. Rehkopf DH, Needham BL, Lin J, et al. Leukocyte Telomere Length in Relation to 17 Biomarkers of Cardiovascular Disease Risk: A Cross-Sectional Study of US Adults. PLoS Med 2016; 13: e1002188.

51. Rehkopf DH, Dow WH, Rosero-Bixby L, Lin J, Epel ES, Blackburn EH. Longer leukocyte telomere length in Costa Rica’s Nicoya Peninsula: a population-based study. Exp Gerontol 2013; 48: 1266–73.

52. Mather KA, Jorm AF, Milburn PJ, Tan X, Easteal S, Christensen H. No associations between telomere length and age-sensitive indicators of physical function in mid and later life. J Gerontol A Biol Sci Med Sci 2010; 65: 792–9.

53. Scheller Madrid A, Rode L, Nordestgaard BG, Bojesen SE. Short Telomere Length and Ischemic Heart Disease: Observational and Genetic Studies in 290 022 Individuals. Clin Chem 2016; 62: 1140–9.

