## Supplemental Materials for "Genetically Increased Telomere Length and Aging-related Physical and Cognitive Traits in the UK Biobank"

**Table of Contents**

**Appendix S1.** Description of simulation parameters to examine selection bias in UKB.

**Figure S1.** Distribution of genetically increased TL (GI-TL) among UKB participants.

**Figure S2.** Causal diagram between telomere length (TL) and aging-related traits in UKB and potential influence of selection

**Figure S3.** QQ-plot of aging-related traits and genetically increased TL (GI-TL) associations stratified by gender.

**Figure S4.** QQ-plot of aging-related traits and genetically increased TL (GI-TL) associations stratified by median age (58 years).

*In separate excel document:*

**Table S1.** Reported associations between genetically increased telomere length (GI-TL) and age-related traits stratified by gender.

**Table S2.** Reported associations between genetically increased telomere length (GI-TL) and age-related traits stratified by median age (58 years).

**Table S3.** Genetically increased TL (GI-TL) and mortality among UK Biobank participants.

**Table S4.** Association between genetically increased telomere length (GI-TL) and aging related traits using population-based inverse probability weights.

**Table S5.** Causal effect estimates per SD increase in telomere length (TL) using Mendelian Randomization (MR) approaches.

**Table S6**. Results from analysis of simulated datasets examining of selection bias under no true effect of risk factor (TL) on outcome (systolic blood pressure [SBP]) (β_x_=0) and selection on outcome (y_y_).

**Table S7.** Results from analysis of simulated datasets examining selection bias under no true effect of risk factor (TL) on outcome (systolic blood pressure [SBP]) (β_x_=0) and selection on exposure (y_x_).

**Table S8.** Results from analysis of simulated datasets examining selection bias under true effect of risk factor (TL) on outcome (systolic blood pressure [SBP]) (β_x_=0.12) and selection on outcome (y_y_).

**Table S9**. Results from analysis of simulated datasets examining selection bias under true effect of risk factor (TL) on outcome (systolic blood pressure [SBP]) (β_x_=0.12) and selection on exposure (y_x_).

**Appendix 1.** Description of simulation parameters to examine selection bias in UKB.

| **Parameter** | **Definition** | **Varying** | **Value** |
| --- | --- | --- | --- |
| α^2^_g_ | PVE by genetic instrument in risk factor | No | 0.04 |
| α^2^_u_ | PVE by confounders in risk factor | No | 0.50 |
| β_x_ | causal effect estimate | No | 0 (none) or 0.12 (observed) |
| β^2^_u_ | PVE by confounders in outcome | No | 0.50 |
| γ_0_ | Prevalence of selection | No | 0.0545 |
| γ_x_ | Risk factor effect on selection | Yes | -2,-1,-0.75,-0.5, 0 ,0.5,0.75,1,2 |
| γ_y_ | Outcome effect on selection | Yes | -2,-1,-0.75,-0.5, 0 ,0.5,0.75,1,2 |
| γ_u_ | Confounder effect on selection | Yes | -1,0,1 |

**Figure S1.** Distribution of genetically increased TL (GI-TL) among UKB participants. Top histogram corresponds to GI-TL, where one unit GI-TL corresponds to one SD increase in TL (~1200 base pairs). Bottom histogram corresponds to GI-TL converted to base pairs.

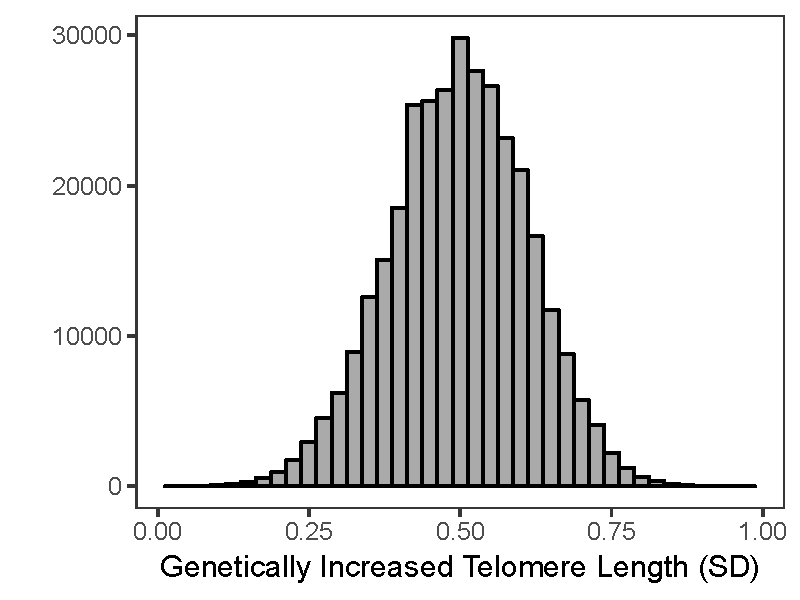

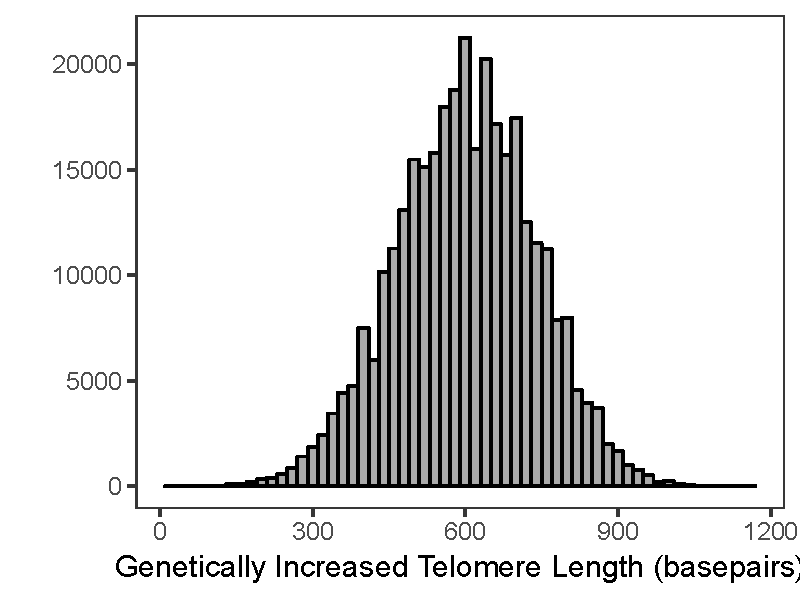

**Figure S2.** Causal diagram between telomere length (TL) and aging-related traits in UKB and potential influence of selection.

Unmeasured confounders

Telomere Length (TL)

Aging-related trait

Genetically Predicted Telomere Length (GP-TL)

Established causal relationship

Selection on exposure (TL)

Selection on outcome (SBP)

Causal relationship to be estimated

Inverse Probability Weighting

Simulation Analysis

**Figure S3.** QQ-plot of aging-related traits and genetically predicted TL (GP-TL) associations stratified by gender. Black and gray dashed lines correspond to Bonferroni corrected significance (p < 0.001) and nominal significance (p < 0.05), respectively.

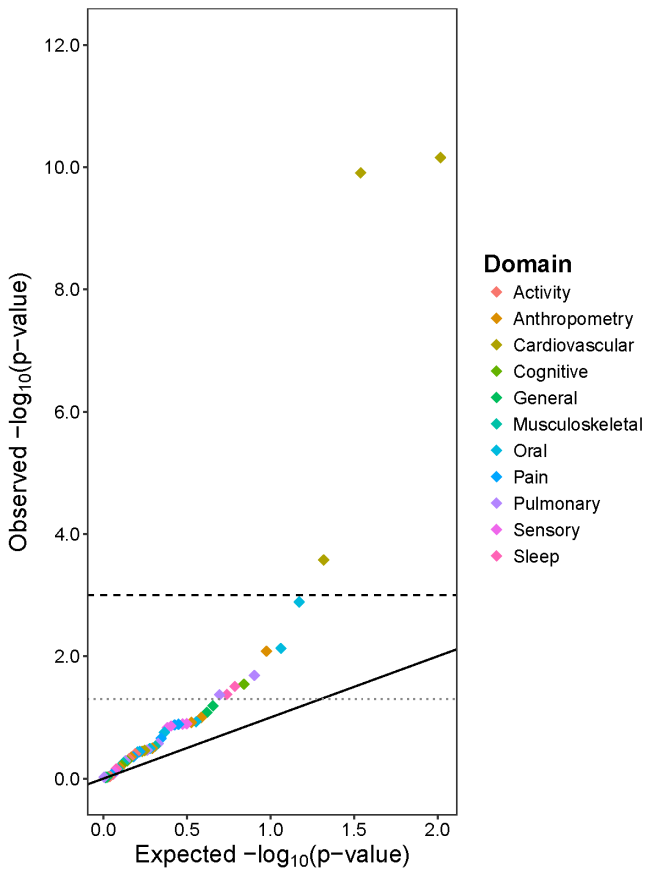

Men (n=156,273)

Women (n=181,272)

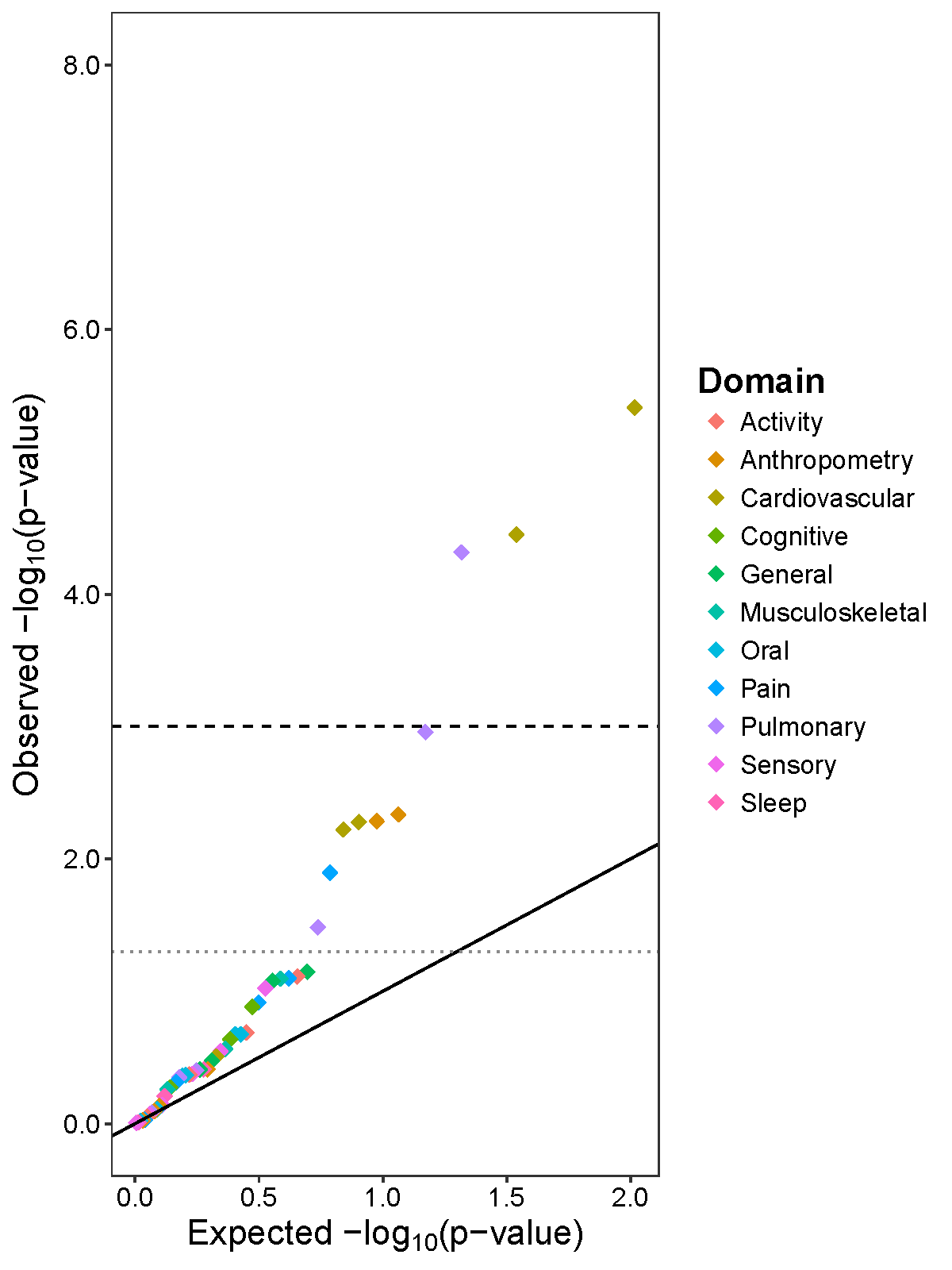

**Figure S4.** QQ-plot of aging-related traits and genetically predicted TL (GP-TL) associations stratified by median age (58 years). Black and gray dashed lines correspond to Bonferroni corrected significance (p < 0.001) and nominal significance p < 0.05, respectively.

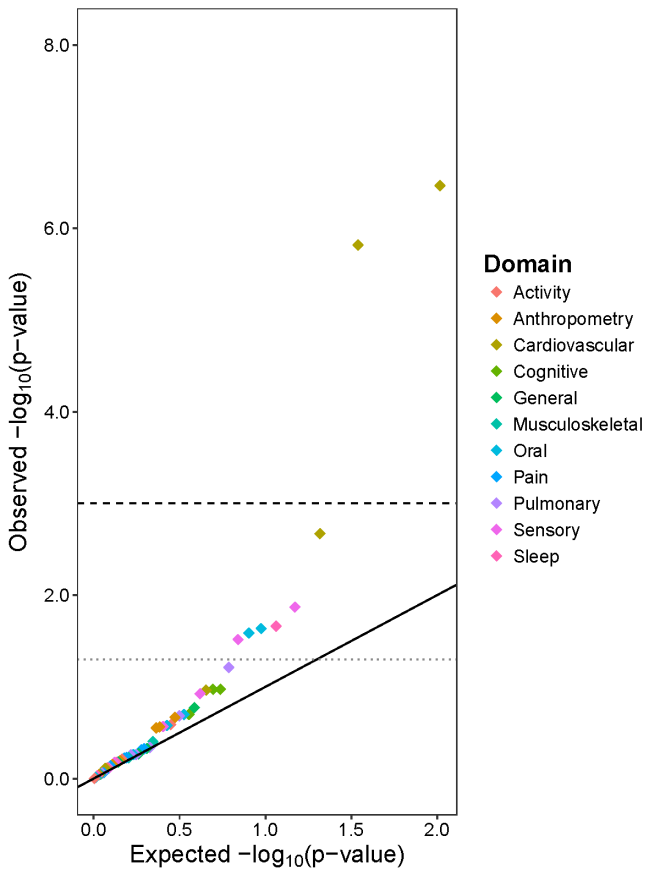

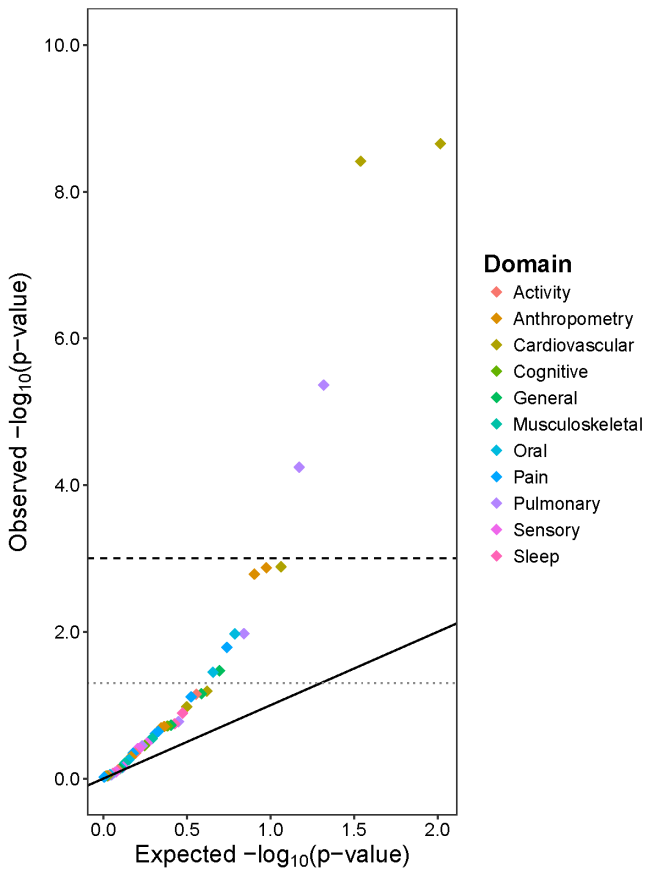

< 58 years (n=158,727)

≥ 58 years (n=178,809)
